# Direct Genome-Scale Screening of *Gluconobacter oxydans* B58 for Rare Earth Element Bioleaching

**DOI:** 10.1101/2024.06.10.598312

**Authors:** Sabrina Marecos, Brooke Pian, Sean A. Medin, Alexa Schmitz, Mingming Wu, J. Brian Balta, Esteban Gazel, Megan Holycross, Matthew C. Reid, Buz Barstow

## Abstract

The transition to a sustainable energy economy will require an enormous increase in the supply of rare earth elements (REE). Bioleaching offers a promising alternative to conventional hydrometallurgical methods for REE extraction from low-grade ores. However, exploiting this potential remains challenging due to large gaps in our understanding of the genetics involved, and inadequate biological tools to address them. We generated a highly non-redundant whole genome knockout collection for the bioleaching microbe *Gluconobacter oxydans* B58, reducing redundancy by 85% compared to the previous best collection. This new collection was directly screened for bioleaching neodymium from a synthetic monazite powder, identifying 89 genes important for bioleaching, 68 of which have not previously been associated with this mechanism. We conducted bench-scale experiments to validate the extraction efficiency of promising strains: 8 demonstrated significant increases in bioleaching by up to 111% (*G. oxydans* δ*GO_1598*, a disruption of the gene encoding the orotate phosphoribosyltransferase enzyme PyrE), and one strain significantly reduced it by 97% (δ*GO_1096*, a disruption of the gene encoding the GTP-binding protein TypA). Notable changes in biolixiviant pH were only observed for 3 strains, suggesting an important role for non-acid mechanisms in bioleaching. These findings provide valuable insights into further enhancing REE-bioleaching by *G. oxydans*’ through targeted genetic engineering.

## Introduction

Rare Earth Elements (REE; including the lanthanides (*Z* = 57 to 71), scandium, and yttrium) are crucial ingredients in present-day sustainable energy technologies including wind turbines^1,2^ and electric vehicles^3^, and in future ones like high-temperature superconductors^4–7^. As a result of this, the demand for REE is expected to grow by between 3- and 7-fold by 2040^8^. However, existing hydrometallurgical methods for REE mining pose considerable environmental risks^9–11^.

The growing demand for an environmentally-friendly supply chain for REE has fueled interest in biotechnological processes for REE mining and recycling^9,12–20^. Bioleaching solubilizes metals from minerals and end-of-life feedstocks with a microbially-secreted mineral-dissolving cocktail called a biolixiviant^13,14,16,21–23^. In recent years, the search for more efficient bioleaching strategies has gained significant interest^23–25^.

The Generally Regarded As Safe (GRAS) acidophilic bacterium *Gluconobacter oxydans*^26–29^ has played a significant role in bioleaching research over the past decade^14,21,30–33^. This microbe offers a unique capability for secretion of organic acids^21,29,34^ and creation of a highly acidic biolixiviant (primarily containing gluconic acid^14^) which is particularly effective for recovering REE from sources including minerals like allanite^15^; industrial residues like phosphogypsum^30^, red mud^35^, and blast furnace slag^36^; and recycled materials like fluid cracking catalyst (FCC)^21^, nickel-metal-hydride (NiMH) batteries^37^, and retorted phosphor powder (RPP)^16,21,35^.

We have already improved bioleaching of REE by *G. oxydans* by first identifying genes involved in acid production with high-throughput screening^16^, and then engineering their regulation^15^. We increased REE-bioleaching from the REE-containing mineral allanite by up to 73% by up-regulating the membrane bound dehydrogenase, *mgdh*, and knocking out the phosphate signaling and transport system *pst*^15^.

Despite progress in enhancing REE-bioleaching with *G. oxydans*, further genetic improvements may be necessary to allow it to leapfrog traditional thermochemical extraction processes^33,38^. One way to further improve it is to target the regulation of additional genes implicated in bioleaching by acid production screens^16^. However, it is proposed that metal solubilization in bioleaching happens through three processes: acidolysis via acid production, complexolysis by creating complexes between metal ions and organic compounds, and redoxolysis, which involves electron transfer and oxidative reactions^33^. *G. oxydans*’ biolixiviant appears to be more effective than pure gluconic acid controls at the same pH^21^. This could suggest that the biolixiviant contains additional components that amplify its effectiveness^21,22,30,32,39^.

We hypothesize that further characterizing the mechanism of bioleaching and then engineering the composition of the biolixiviant is the route to future significant increases in bioleaching efficiency. But the acid production screens that we have used to date may not be sensitive to complexolysis and redoxolysis. Thus, to fully characterize the biolixiviant, it is necessary to identify genes directly linked to bioleaching, rather than just acid production alone. In this study, we conducted a genome-scale screen of a new quality-controlled *G. oxydans* B58 whole genome knockout collection with a high-throughput assay designed to directly identify genes involved in REE-bioleaching.

## Results

### *G. oxydans*’ Quality-Controlled Collection Is 85% Smaller Than Previous Collection

To facilitate direct high-throughput screening of mineral-dissolution, we built a highly non-redundant Quality-Controlled (QC) whole genome knockout collection for *G. oxydans* B58. The QC collection was built by removing redundancy from our earlier condensed collection (CC) of *G. oxydans* transposon disruption mutants^16^. Throughout this article, we refer to transposon disruption mutants with a ‘δ’ symbol (as opposed to a ‘Δ’ symbol for clean deletion mutants; *e.g*., δ*GO_1096*).

In total, the QC collection contains a single carefully chosen transposon disruption mutant for 2,733 non-essential genes in the *G. oxydans* B58 genome arrayed onto 57 checker-boarded 96-well plates (**Supplementary Data S1**). The QC collection is 94% smaller than our *G. oxydans* saturating-coverage transposon mutant collection used to source all mutants in our earlier work (containing 49,256 mutants)^16^, and 85% smaller than our previous best curated collection (17,706 mutants)^16^. An overview of QC collection construction is shown in **Figure 1**, and collection statistics are shown in **Table 1**. Full details of QC collection construction can be found in **Materials and Methods**.

**Figure 1.**
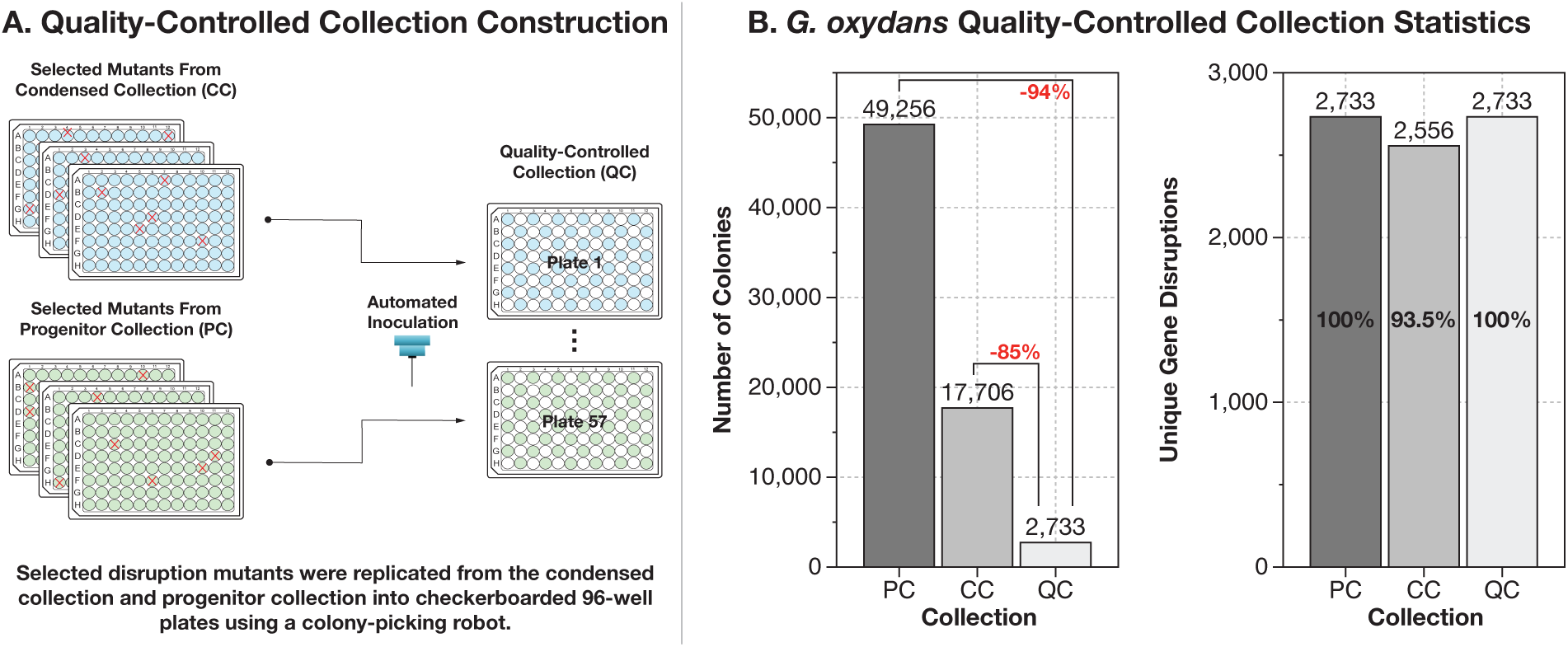
Construction of a quality-controlled (QC) collection containing a single disruption mutant for 2,733 genes in *Gluconobacter oxydans*’ genome. (**A**) The QC collection was built by choosing the best available mutants from an earlier condensed collection (CC) of *G. oxydans* mutants^16^, and by supplementation from the originating saturating-coverage transposon collection (Progenitor Collection or PC)^16^. Mutants were transferred from the earlier collections by fluid transfer into fresh 96-well plates with a colony picking robot. (**B**) The QC collection contains 2,733 unique mutants distributed across 57 checker-boarded 96-well plates. This collection represents a 94% reduction in size from the PC, and an 85% reduction in size from the CC, indicated in red. It is worth noting, that the CC collection contained disruption mutants with confirmed identity for 2,556 genes (93.5% of the 2,733 genes disrupted in the PC), whereas the QC collection aims to include a disruption mutant for 100% of genes disrupted in the PC.

**Table 1.** Overview of *Gluconobacter oxydans* B58 quality-controlled (QC) whole genome knockout collection statistics. This table summarizes the transposon location characteristics of the mutants derived from the progenitor collection (PC) and condensed collection (CC) used to generate the QC collection. Mutants were categorized based on the distance of the disruption from the translation start of the gene: less than 50 base pairs, at least 50 base pairs, within 0.5 fractional distance of the gene, beyond 0.5 fractional distance, and disruptions located at least 50 base pairs downstream and within 0.5 fractional distance, which were considered the optimal locations for disruption of gene function.

| Collection | Number of Mutants | Distance from Translation Start |  |  |  |  |
| --- | --- | --- | --- | --- | --- | --- |
|  |  | < 50bp | ≥ 50bp | ≤ 0.5 | > 0.5 | ≥ 50 bp AND ≤ 0.5 |
| Progenitor Collection (PC) | 285 | 21 | 264 | 172 | 113 | 153 |
| Condensed Collection (CC) | 2,448 | 240 | 2,208 | 2,037 | 411 | 1,797 |
| Total (QC Collection) | 2,733 | 261 | 2,472 | 2,209 | 524 | 1,950 |

### Genome-Scale Screen of QC Collection Identifies 68 Genes Not Previously Implicated in REE-Bioleaching

We performed a genome-scale screen of the QC collection to identify disruption mutants with differential REE-bioleaching capabilities. We developed a rapid microplate-based colorimetric assay with the REE-chelating dye Arsenazo-III (As-III) to screen for REE extraction^18,40,41^ from a synthetic monazite powder (**Figures 2A** and **2B**) (monazite is the mineral host for REE at the Mountain Pass Mine in California, one of the largest REE mines in the world^10^). Unlike natural monazite (which contains multiple light lanthanides), this synthetic mineral contains just neodymium^42^, allowing greater consistency in the determination of extraction efficiency. Two synthetic monazite powder batches were synthesized (Mon1 and Mon2) as described by Balta *et al.*^42^, presenting minor differences in Nd-phosphate composition (**Supplementary Data S2**). Mon1 was composed of monazite-Nd (NdPO4), rhabdophane-Nd (NdPO4·2H2O), and neodymium oxyphosphate (Nd3PO7). In contrast, Mon2 consisted solely of monazite-Nd and rhabdophane-Nd.

**Figure 2.**
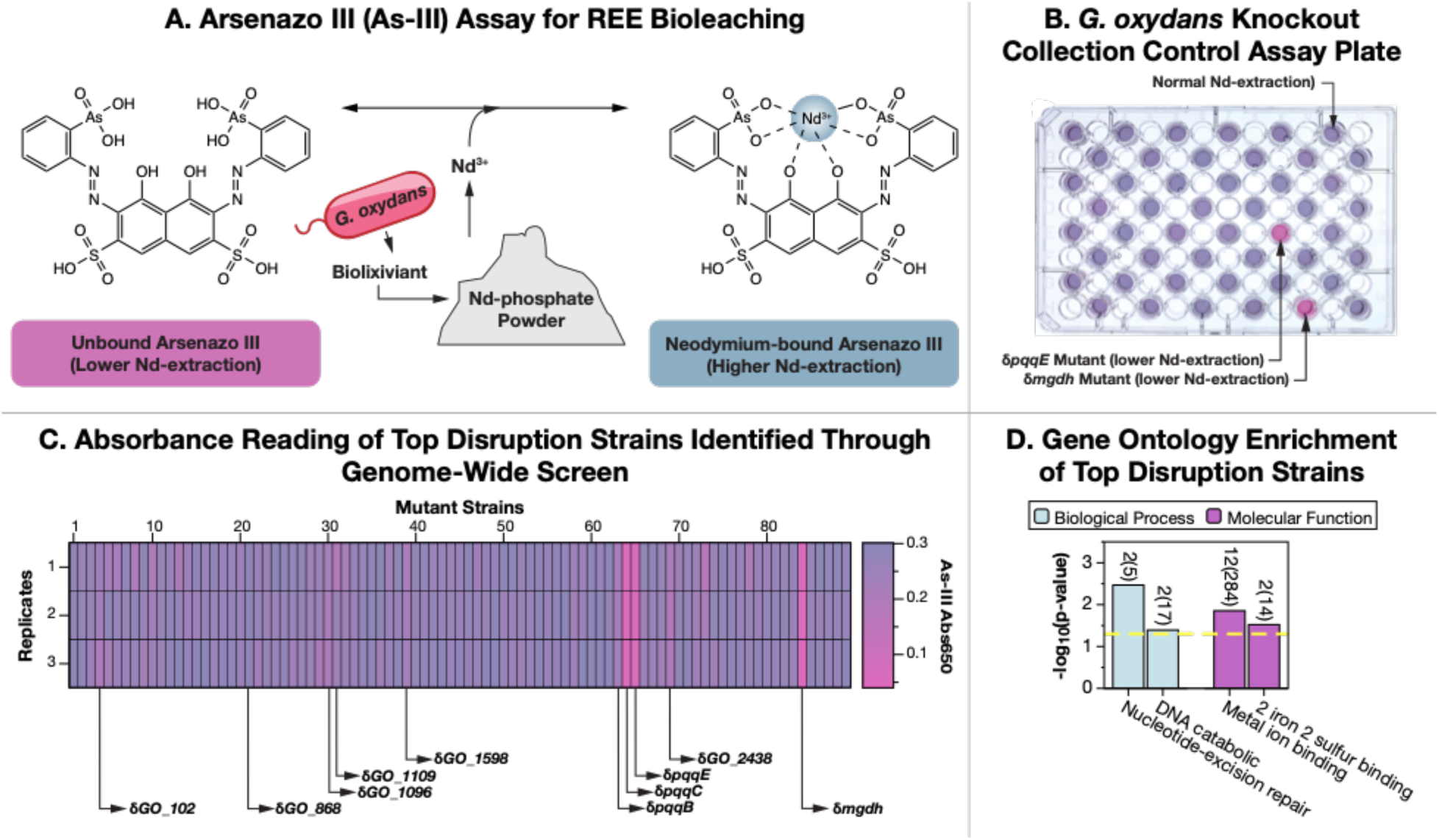
Direct bioleaching screening of the quality-controlled whole genome knockout collection of *Gluconobacter oxydans* identifies 68 gene disruptions not previously associated with bioleaching. (**A**) We developed a rapid 96-well plate based high-throughput screen for REE-bioleaching that uses a colorimetric assay with the Arsenazo III (As-III) REE-chelating dye in presence of synthetic monazite. This assay directly detects REE release, facilitating the identification of disruptions with differential bioleaching capabilities. (**B**) Color differences in each well identify which genes control bioleaching. Gene knockouts that reduce bioleaching leave the As-III dye unbound and pink, while knockouts that increase bioleaching turn the dye blue. (**C**) The genome-scale screening of the QC collection identified 89 hits including 68 gene disruptions not previously associated with the bioleaching capabilities of *G. oxydans.* (**D**) Gene Ontology (GO) Enrichment was performed with a Fisher’s Exact Test (*p <* 0.05, yellow dashed line) to analyze the hits identified. Significantly enriched GO terms with at least 2 representatives are displayed, and numbers above the bars represent the significant hits out of the total annotated genes associated with that gene ontology in the genome of *G. oxydans* (the full list of GO terms can be found in **Supplementary Data S5**). Only terms belonging to the Biological Process and Metabolic Function GO groups were significantly enriched.

Mon1 demonstrated greater dissolution than Mon2 during bioleaching. This discrepancy is presumed to be due to the elevated concentration of neodymium oxides in Mon1 (**Supplementary Data S2**), along with the higher thermal expansion and oxidation state of the Nd3PO7 ^43,44^. We utilized both Mon1 and Mon2 in our screening to leverage their distinct characteristics. Mon1 enhanced the detection sensitivity for potential hits in the initial preliminary survey, thereby expanding the selection pool for subsequent validation. Mon2, being less reactive, allowed for the isolation of the most significant outliers in the initial survey, offering a closer analog to bioleaching with natural monazite.

Our preliminary survey (**Supplementary Data S3**) identified 89 gene disruptions that produced significant changes in Nd extraction from the Mon1 synthetic monazite (**Figure 2C**). These hits included disruption of 21 genes previously linked to acidification^16^, and 68 genes that had not previously been linked to bioleaching (**Supplementary Data S4**).

A follow-up assay of these 89 mutants found 6 mutants that produced statistically significant changes to Nd-extraction from the Mon2 synthetic monazite (**Supplementary Data S5**, **Table 2**). We added an additional 6 mutants for follow up study that were prominent in the initial Mon1 Nd-extraction survey (**Table 2**).

**Table 2.** Selected gene disruptions for direct measurement of bioleaching. This table lists the selected gene disruption mutants for analysis, specifying their gene identifiers and disrupted features. It also presents the percentage change in extraction efficiency compared to pWT, where positive values indicate enhanced bioleaching, and negative values denote a reduction.

| Gene Identifier | Feature Disrupted | Extraction<br>Against pWT (%) |
| --- | --- | --- |
| GO_1096 | GTP-binding protein TypA/BipA | -97 |
| GO_1598 | Orotate phosphoribosyltransferase (EC 2.4.2.10) | +111 |
| GO_102 | hypothetical protein | +102 |
| GO_1109 | Transcriptional regulator, LysR family | +89 |
| GO_1113 | Xanthine and CO dehydrogenases maturation factor, XdhC/CoxF family | +12 |
| GO_2555 | hypothetical protein | +74 |
| GO_2946 | hypothetical protein | +91 |
| GO_1162 | Phosphate regulon transcriptional regulatory protein PhoB (SphR) | +11 |
| GO_19 | TonB-dependent receptor | +82 |
| GO_1968 | Formamidopyrimidine-DNA glycosylase (EC 3.2.2.23) | +17 |
| GO_1991 | Efflux ABC transporter for glutathione/L-cysteine, CydC subunit | +56 |
| GO_1816 | Transcriptional regulator, YafY family | +83 |

### Metal Binding and DNA Repair Gene Ontologies are Highly Enriched in Bioleaching Hits

The 68 genes not previously associated with REE-bioleaching that were identified through the initial Mon1 REE-extraction screen were analyzed for gene ontology (GO) enrichment with topGO^45^. A total of four GO terms were found to be significantly enriched (*p* < 0.05 by Fisher’s Exact Test), and to be associated with more than one gene (**Supplementary Data S6**). These four enriched GO terms belong to the biological process (BP) and molecular function (MF) groups (**Supplementary Data S6**). No significant terms belonging to the cellular component group were found.

The two most statistically-significantly enriched GO terms are metal ion binding and nucleotide-excision repair (**Figure 2D**, **Supplementary Data S6**). We speculate that disruption of these genes affects bioleaching indirectly by modifying the expression of metabolic pathways that produce metal solubilizing compounds. For example, metal ion binding proteins are often associated with signaling, sensing, and regulation^46–49^, suggesting that their disruption could alter *G. oxydans*’ bioleaching capabilities. Similarly, disruption of the nucleotide-repair system could impair the ability of *G. oxydans* to respond to damage generated by oxidative stress resulting from initial biolixiviant production^50,51^ and thus alter further biolixiviant production.

### Gene Disruptions of Interest Demonstrate Significant Changes in Bioleaching

The 12 gene disruption mutants that were highlighted by the Mon1 and Mon2 high-throughput assays (**Table 2**) were validated with bench-scale experiments where bioleaching was measured by Inductively Coupled Plasma - Mass Spectrometry (ICP-MS) (**Figure 3**). When compared to the proxy wild-type strain (pWT) (See “**Materials and Methods**”), 9 of the 12 disruption mutants generated significant changes in bioleaching (*p* < 0.05). Eight disruption mutants produced greater (with increases of between 56 and 111%) Nd-bioleaching than pWT, while only one mutant (δ*GO_1096*) produced a reduction (-97%) in bioleaching (**Figure 3**). When compared to the wild-type strain (WT), all 12 disruption mutants tested produced significantly different extractions (*p* < 0.01) (**Supplementary Figure S1**), showing increases from 75 to 231%, and a 95% reduction by δ*GO_1096*.

**Figure 3:**
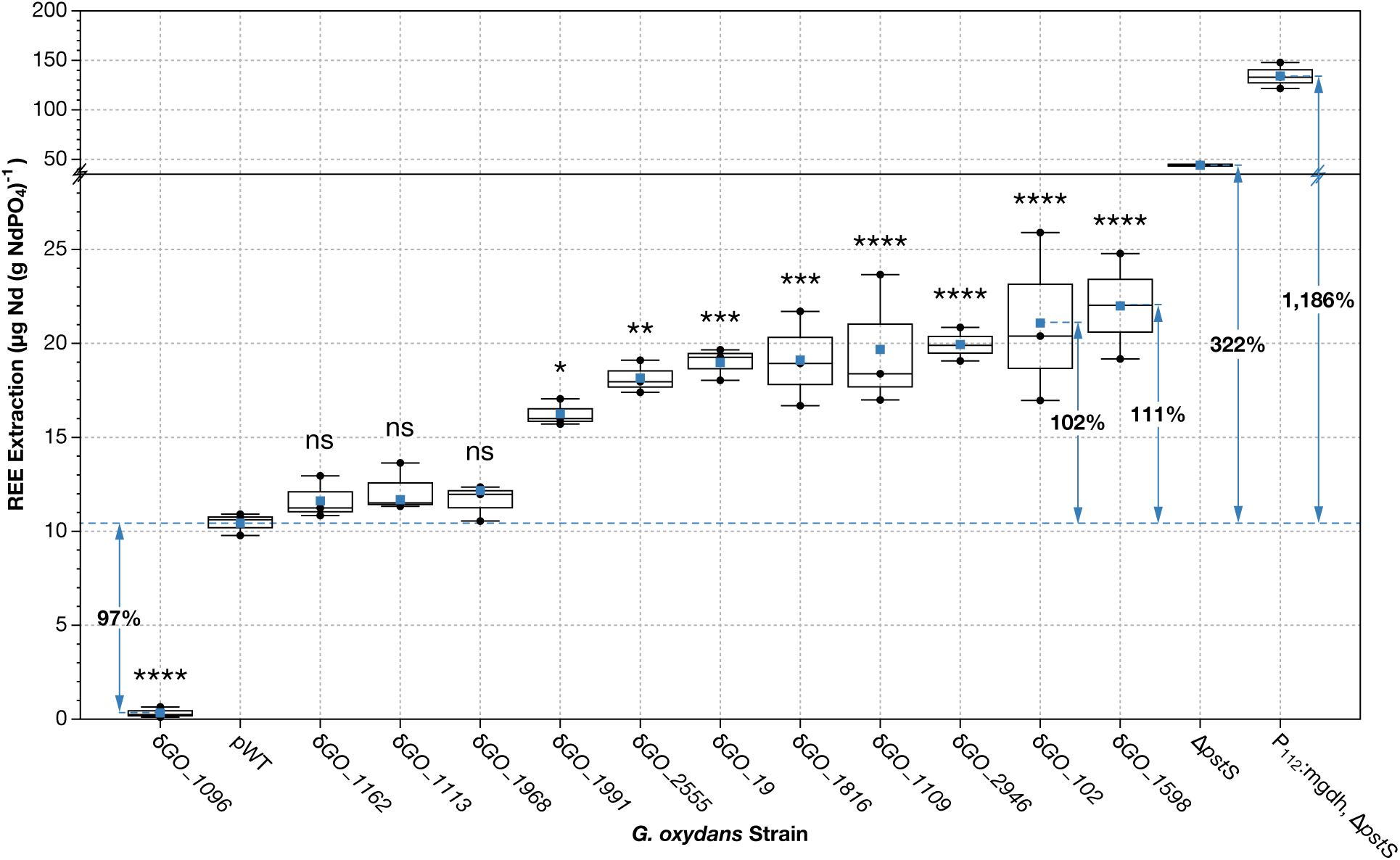
Identified *Gluconobacter oxydans* gene disruptions raised REE-bioleaching by up to 111% or lowered it by 97%. Direct measurement of bioleaching extraction was performed on disruption mutants of interest following bioleaching experiments with synthetic monazite, using ICP-MS analysis. REE extractions were analyzed with pairwise comparisons among disruption mutants (*n* = 3) and pWT (*n* = 3). Levels of extraction significantly different from pWT are labeled with asterisks (*\*p <* 0.05*; **p <* 0.01*; ***p <* 0.001*, ****p <* 0.0001) or “ns” if *p* > 0.05, denoting statistical significance after Bonferroni-corrected *t*-tests (*N* = 12). Blue squares indicate the mean extraction for each mutant, the center line denotes the median, boxes show the upper and lower quartiles, and whiskers extend to the range of data points within 1.5 times the interquartile range. Strains δ*GO_1162*, δ*GO_1113*, and δ*GO_1968* did not show statistically-significant changes in extraction compared to pWT. Only one disruption mutant, δ*GO_1096*, decreased bioleaching, demonstrating a 97% reduction in extraction. Current best-performing engineered strains, *G. oxydans* Δ*pstS* and *G. oxydans* P112:*mgdh*, Δ*pstS*, were included to contrast the extractions of the tested disruption mutants. Significant changes in extraction efficiency compared against the WT strain can be found in **Supplementary Figure S1**.

Even though the increases in Nd-bioleaching created by 9 of the newly identified gene disruptions are large, they are notably smaller than those produced by our current two best-performing strains (*G. oxydans* Δ*pstS* and *G. oxydans* P112:*mgdh*, Δ*pstS*^15^; demonstrating 322% and 1,186% increase over pWT (**Figure 3**), and 564% and 1,922% over WT (**Supplementary Figure S1**).

### Many Gene Disruptions of Interest Do Not Affect Biolixiviant pH

The pH of the biolixiviants produced by the 12 gene disruption mutants selected was measured (**Figure 4**). Unsurprisingly, given our focus on identifying mutants absent in acid-production screens, yet still noteworthy, 9 of the 12 mutants did not exhibit significant changes in biolixiviant pH when compared to a proxy wild-type strain of *G. oxydans*. Only the δ*GO_102*, δ*GO_1096*, and δ*GO_1109* mutants produced statistically-significant changes in biolixiviant pH (**Figure 4**). δ*GO_102* and δ*GO_1109* produced very modest reductions in biolixiviant pH of 0.06 and 0.07 units respectively, which are only barely above the precision-threshold of a pH meter. δ*GO_1096* produced a large increase in biolixiviant pH of 0.55 units.

**Figure 4:**
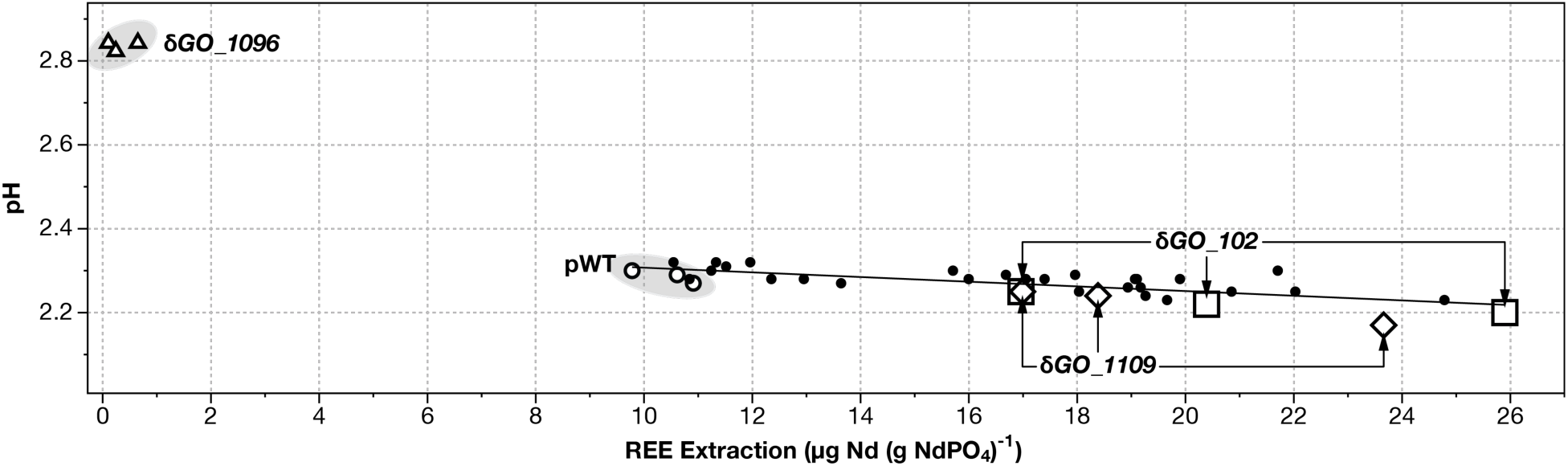
Only three of the identified *Gluconobacter oxydans* gene disruptions produce statistically-significant changes in biolixiviant pH. REE extraction and biolixiviant pH were plotted for each strain selected for direct bioleaching measurement. Strain replicates that significantly changed the biolixiviant pH compared to proxy wild-type are indicated as follows, pWT “○”, δ*GO_102* “□”, δ*GO_1109* “◇”, and δ*GO_1096* “Δ”. Other gene disruptions with data points falling within the model are represented as “●”, including data points for δ*GO_19*, δ*GO_1113*, δ*GO_1162*, δ*GO_1598*, δ*GO_1816*, δ*GO_1968*, δ*GO_1991*, δ*GO_2555*, and δ*GO_2946*.

## Discussion

The generation of the QC collection facilitates the implementation of cost-effective and elaborate screening methods^52,53^. The curated collection encompasses disruption mutants for 2,733 genes, one single representative for almost every non-essential gene in *G. oxydans*’ genome^16^. The reduction in redundancy is crucial to improve the utility of a knockout collection, improving its value as a practical tool for the exploration of gene function^53,54^. This streamlined collection was essential for conducting a technically challenging genome-wide screen for REE-bioleaching, leading to the identification of previously uncharacterized aspects of this mechanism. To our knowledge, this study represents the first direct whole genome screening for bioleaching of any mined metal.

In total, we identified 89 gene disruptions involved in Nd-bioleaching from synthetic monazite. Notably, 21 genes implicated in acidification, previously reported^16^, were among the hits found through this screen, corroborating their direct involvement in REE-bioleaching. This outcome verifies the reliability of our As-III screen for Nd-bioleaching but also underscores some challenges related to reduced sensitivity. However, we also identified an additional 68 gene disruptions that were tested in our earlier assays but were not implicated in bioleaching. Out of 12 disruption mutants selected for further study, 9 did not produce any statistically-significant change in pH (**Figure 4**). This indicates that the new screen is more comprehensive than pH-indicator screens for acid production. Furthermore, these results suggest that additional components might be present during bioleaching by *G. oxydans*^13,30^, potentially increasing solubilization.

Notable and significant changes in bioleaching were validated by ICP-MS through bench-scale experiments (**Figures 3** and **4**). Out of the 12 mutants selected for verification, only one (δ*GO_1096*) reduced extraction, suggesting that this gene normally facilitates bioleaching. In contrast, 11 of the mutants appear to regulate bioleaching, and their disruption removes the inhibitory controls on it.

Three results (δ*GO_1096*, δ*GO_1991*, and δ*GO_1968*) imply that changes in stress response pathways can both improve or impair the bioleaching mechanism. This notion has been discussed by Sousa *et al*. after observing increased resistance to acetic acid at specific levels of oxidative stress in *Saccharomyces cerevisiae*^55^.

Disruption of the gene encoding Formamidopyrimidine-DNA Glycosylase Fpg (*GO_1968*; belonging to both the metal ion binding and nucleotide excision repair gene ontologies terms) increased bioleaching. Fpg is a glycosylase involved in the repair of DNA damage caused by oxidative stress^56^, and its disruption might lead to a heightened cellular response. For instance, it has been shown that certain levels of mistranslation can improve *Escherichia coli*’s tolerance to oxidative stress^51^. The metabolic response to accumulated DNA damage could involve transcription errors and shifts in secondary metabolite production. Such shifts might influence the types or quantities of solubilizing compounds produced by *G. oxydans*. Moreover, the gene coding for the Efflux ABC Transporter for Glutathione CydC (*GO_1991*) plays a role in redox balance within the periplasm during oxidative stress conditions^57^, and its disruption also increases bioleaching.

Notably, only a single gene disruption led to a decrease in bioleaching efficiency (δ*GO_1096*). *GO_1096* codes for the GTP-binding protein TypA which has been linked to stress response^58^. This suggests that its disruption might hamper *G. oxydans*’ ability to adapt to the stress generated during bioleaching, thereby diminishing efficiency. δ*GO_1096* was the only mutant that increased biolixiviant pH (out of only 3 that changed the pH at all). This result suggests TypA may facilitate acid-mediated bioleaching as well as non-acid bioleaching. We suspect that this gene was not found in our earlier acid-production screen^16^ because we swapped the *GO_1096* disruption mutant used in our original curated collection^16^ for a new choice that was more likely to disrupt gene function. Interestingly, δ*GO_1096*’s increase in biolixiviant pH is modest compared to those produced by mutants identified in our previous screen that produced similar reductions in REE-extraction. For example, disruption of the *tldD* gene, necessary for the synthesis of the PQQ molecule (a cofactor for the membrane bound glucose dehydrogenase), produced a 92% reduction in REE-extraction (admittedly from a different substrate) but raised biolixiviant pH by ≈ 0.7 units^16^.

Furthermore, we observed that the disruption of three genes involved in gene expression increased bioleaching: *GO_1162* encoding the phosphate regulon regulatory protein PhoB, *GO_1109* encoding a LysR substrate binding domain, and *GO_1816* encoding the transcriptional regulator YafY.

PhoB is involved in the regulation of inorganic phosphate (Pi) homeostasis^59^; it can positively or negatively regulate phosphate uptake in the presence of excess or limited Pi^60^. Disrupting this transcriptional activator causes cells to lose the ability to modulate the phosphate regulon, which proves to be advantageous during bioleaching in phosphate-rich environments.

The LysR substrate binding domain binds specific ligands for signal recognition, and as a result, triggers the activity of transcriptional regulators^61^. Its disruption may influence gene repression or activation, altering the metabolic flux in a manner favorable for bioleaching. The regulatory protein YafY presents HTH (helix-turn-helix) and WYL (tryptophan, W; tyrosine, Y; and leucine, L) domains, which are known for their role in gene expression in response to DNA damage^62^. Disruption of the hypothetical protein GO_102 produced the second largest increase in Nd-bioleaching (**Figure 3**). *blastp* analysis^63^ revealed 100% sequence alignment between GO_102 and an HTH-domain containing protein, suggesting it might also be involved in gene expression. These disruptions may lead to significant changes in gene expression levels, which could explain the increase in extraction.

Both δ*GO_102* and δ*GO_1109* disruption mutants decreased biolixiviant pH but were not identified by our earlier acid production screen. The pH changes caused by these mutants are modest compared with prominent mutants from our earlier acid production screen (which decreased pH by 0.1 to 0.2 units)^16^. We suspect that these pH changes were too small to be detected by our earlier screen. This leads us to presume that these mutants may be simultaneously altering acid and non-acid mechanisms of bioleaching.

We speculate that disruption of the hypothetical protein GO_2946 might enhance bioleaching by increasing the permeability of the outer membrane of *G. oxydans*. *blastp* analysis^63^ showed homology of GO_2946 to a glycosyltransferase family 2 (GT2) protein, whose function involves the transfer of sugar moieties among intermediates during the biosynthesis of polysaccharides^64^. In a previous study, a GT2 protein was found to repress oxidative fermentation in *Gluconacetobacter intermedius*; and when disrupted, final yields of gluconic acid and acetic acid increased compared to the parental strain^64^. Changes in the polysaccharide composition of the cell surface might increase the permeability of the membrane, facilitating the diffusion of substrate, therefore enhancing transport of the biolixiviant out of the cell^64^.

The disruption of a TonB-dependent receptor gene (*GO_19*) increased bioleaching. This type of system is often associated with the uptake of iron bound to siderophores^65^. Under iron starvation conditions, bacterial cells tend to up-regulate secretory mechanisms in an attempt to manage iron deficiency^66,67^. Disruption of this gene might alter iron sensing and transport, prompting the cell to increase biolixiviant production.

Disruption of genes involved in nucleotide metabolism increased bioleaching (*GO_1598, GO_1113,* and *GO_2555)*. The largest increase in bioleaching was produced by δ*GO_1598*. *GO_1598* codes for the orotate phosphoribosyltransferase enzyme PyrE which catalyzes the production of orotidine monophosphate (OMP), using phosphoribosyl diphosphate (PRPP) and orotate^68^. One way that the disruption of *GO_1598* could increase bioleaching is by increasing availability of orotate, which is known to chelate metals^68^, and in particular lanthanides^69^. *GO_1113* codes for XdhC, an accessory protein involved in purine metabolism that binds to Molybdenum Cofactor (MoCo) for maturation of xanthine dehydrogenase^70^. We speculate that its disruption could create an accumulation of MoCo, and accelerate other enzymatic reactions used in the production of the biolixiviant. *GO_2555* codes a hypothetical protein that demonstrated partial homology^63^ (E-value = 1×10^−125^) to the L-dopachrome tautomerase yellow-f2 protein (EC 5.3.2.3) involved in tyrosine metabolism that interacts with quinone and flavonoids to produce melanin^71–73^. The enhanced bioleaching observed from this disruption may be attributed to the accumulation of intermediates, which can be utilized in other metabolic pathways associated with bioleaching.

Our results propose that the choice of bioleaching substrate can dramatically amplify (or perhaps negate) the effectiveness of genetic engineering. In our earlier work on bioleaching from retorted phosphor powder with *G. oxydans*^16^, the largest genetically-caused increase in bioleaching was 18% (due to disruption of *pstC*). Likewise, an engineered double mutant of *G. oxydans* (with clean deletion of *pstS*, and up-regulation of *mgdh* with the P112 promoter) improved bioleaching from the mineral allanite by up to 73%^15^. In this work on bioleaching monazite, single gene disruptions can improve bioleaching by up to 111%, while the engineered double mutant improves bioleaching by 1,186% over proxy wild-type. It is easy to infer why the effectiveness of the *G. oxydans* biolixiviant could vary from substrate to substrate, as different minerals will have different chemistries that will be affected more or less by the bioleaching mechanisms. However, it is both more challenging and intriguing to explain why the same genetic interventions would produce different relative changes in bioleaching performance.

It remains unclear whether modification of the genes discussed in this article could lead to transformative advancements in bioleaching by *G. oxydans*. It is worth noting that when we first observed the effects generated by transposon disrupted genes, they were very modest compared with their subsequent genetic modifications. For example, disruption of *pstC* only increased bioleaching from retorted phosphor powder by 18% (and disruption of *pstS* did not significantly improve it at all)^16^, but clean deletion of *pstS* improved bioleaching from allanite by 30%^15^. Remarkably, combination of this knockout and up-regulation of *mgdh* increased bioleaching from allanite by 73%^15^. This suggests (and it is emphasized that further testing is necessary) that multiple modifications inspired by the results presented here could have very big impacts on bioleaching performance from some substrates.

## Conclusions

Our work pivots away from the well-characterized acidolysis mechanism to examine contributions to bioleaching via alternative routes. Our findings introduce additional genes that notably influence bioleaching yet remained undetected in acidification screens (**Figures 2** and **3**). Most genes that we identified do not significantly affect biolixiviant pH (**Figure 4**).

These results, along with earlier observations by Reed *et al.*^21^, strongly suggest that the bioleaching mechanism of *G. oxydans* is not purely acid-based. Our data indicates that disrupting genes involved in nucleotide synthesis, transcriptional regulation, DNA repair, nutrient uptake, and oxidative stress response could significantly impact bioleaching. But there is no clear evidence in this list that implies a single gene coding for a redox molecule (*e.g.*, a flavin) or chelator (*e.g.*, a siderophore) amplifies the effectiveness of bioleaching.

All of the gene disruptions we have identified appear to be involved in regulatory functions. This suggests that if there are genes involved in complexolysis or redoxolysis within the *G. oxydans* genome, they could be multiply-redundant, and that knocking out a single gene contributes only marginally to changes in bioleaching. Alternatively, these genes may possess some degree of essentiality, rendering their identification challenging. This highlights the need for broader and higher-sensitivity screening approaches that can capture a more diverse array of bioleaching-related genetic activities, potentially leading to further enhanced methods for harnessing and optimizing the bioleaching mechanism.

## End Notes

### Data Availability

The datasets generated and analyzed during the current study are available from the corresponding author (B.B.).

### Code Availability

Code used in the generation of figures and analysis of data for this article is available from the corresponding author (B.B.).

### Materials and Correspondence

Correspondence and material requests should be addressed to B.B. Individual strains (up to ≈10 at a time) are available at no charge for academic researchers. We are happy to supply a duplicate of the entire *G. oxydans* knockout collection to academic researchers but will require reimbursement for materials, supplies, and labor costs. Commercial researchers should contact Cornell Technology Licensing for licensing details.

## Acknowledgments

This work was supported by Cornell University startup funds, an Academic Venture Fund award from the Atkinson Center for Sustainability at Cornell University, a Career Award at the Scientific Interface from the Burroughs Welcome Fund to B.B., ARPA-E award DE-AR0001341 to B.B., M.H., E.G., and M.W., and a gift from Mary Fernando Conrad and Tony Conrad to B.B. S. Marecos. was supported by a Link Foundation Graduate Fellowship. A.S. was supported by a Cornell Energy Systems Institute Postdoctoral Fellowship. S.A.M. was supported by a Cornell Presidential Life Sciences Graduate Fellowship.

## Contributions

Conceptualization, S. Marecos. and B.B.; Methodology, S. Marecos. and B.B.; Investigation, S. Marecos., B.P., S.A.M., and A.S.; Writing—Original draft, S. Marecos. and B.B.; Writing—Review and editing, S. Marecos., B.B., J.B.B., M.W., E.G., S.A.M., and A.S.; Funding acquisition, S. Marecos., S.A.M., A.S., M.W., M.H., E.G., and B.B.; Resources, M.C.R., E.G., J.B.B., and B.B.; Supervision, M.W., M.H., E.G., M.C.R., and B.B.; Data curation, S. Marecos. and B.B.; Visualization, S. Marecos. and B.B.; Formal analysis, S. Marecos.

## Materials and Methods

### Curation of the Quality-Controlled Whole Genome Knockout Collection

The QC collection was built by removing redundancy from an earlier condensed collection (CC) of *G. oxydans* transposon mutants and supplementing with additional mutants from a saturating-coverage progenitor collection (PC) from the same work^16^. The progenitor collection of transposon mutants was catalogued using the Knockout Sudoku probabilistic inference combinatorial pooling method^52,74^.

The best-choice disruption mutant for each gene in the QC collection was selected by considering the location of the transposon disruption in the gene, and the reliability of the location assignment of the mutant. We followed the generally-accepted consensus that the best location for a transposon to disrupt gene function is between 50 bp from the start of the coding region and the middle of the gene (a fractional distance of ≤ 0.5 into the gene). Furthermore, we preferred to select mutants which location was directly inferred by the Knockout Sudoku algorithm (also known as unambiguous location inference^52,74^), rather than relying upon probabilistic location inference (also known as ambiguous location inference^52,74^). If given the choice between a mutant with non-ideal locations and unambiguous location inference, and a mutant in the same gene with ideal location and ambiguous location inference we chose the former. Mutant genomic location statistics are shown in **Table 1**. A catalog of the QC collection is included in **Supplementary Data S1**.

Mutants were transferred from the earlier collections by fluid transfer with a colony picking robot (CP7200, Norgren Systems, Ronceverte, West Virginia, USA) into fresh 96-well polypropylene storage plates containing yeast peptone mannitol (YPM) media (5 g L^-1^ yeast extract (catalog number C7341, Hardy Diagnostics, Santa Maria, California, USA); 3 g L^-1^ peptone (catalog number 211677, BD, Franklin Lakes, New Jersey, USA); 25 g L^-1^ mannitol (catalog number BDH9248, VWR Chemicals, Radnor, Pennsylvania, USA)) and 100 μg mL^-1^ kanamycin. Plates were incubated at 30 °C and 800 rpm in a Multitron shaker (Infors Inc., Laurel, Maryland, USA) for 3 nights to achieve saturation. Following incubation, stocks were prepared by adding glycerol (catalog number G5516, Sigma-Aldrich, St. Louis, Missouri, USA) to 20% (v/v) to each well for long term storage at -80 °C.

### Genome-Scale REE-Bioleaching Screen

REE concentrations were measured using the REE-chelating Dye Arsenazo-III (As-III; catalog number 110107, Sigma-Aldrich, St. Louis, Missouri, USA). The As-III solution was prepared following the method described by Hogendoorn *et al*.^40^.

The high-throughput screen consisted of a preliminary survey carried out in duplicate followed by a validation screen performed in triplicate. It spanned several weeks and was partitioned into batches of 10-15 plates. Plates containing mutants to be analyzed were replicated from the QC collection using a 96-pin cryo-replicator press (catalog number CR1100, EnzyScreen, Heemstede, Netherlands) into 96-well polypropylene round-bottom plates filled with 150 µL YPM supplemented with 100 µg mL^-1^ kanamycin.

Replicated plates were incubated for two nights at 30 °C, 80% humidity, and 800 rpm in a Multitron shaker (Infors Inc., Laurel, Maryland, USA). Then, cultures were back-diluted either in duplicate or triplicate, by inoculating 5 µL of bacterial culture in 120 µL of the same medium using the PLATEMASTER pipetting system (catalog number F110761, Gilson Inc., Middleton, Wisconsin, USA). After two nights of incubation, the optical density was measured using a plate reader (catalog number 18531, Biotek, Charlotte, Vermont, USA), and the biolixiviant production was set up by measuring the remaining volume in each well and adding glucose (catalog number 0188, VWR Chemicals, Radnor, Pennsylvania, USA) to 20% w/v. Plates were incubated for 24 hours at 30 °C, 80% humidity, and 800 rpm. For all the aforementioned steps, 96-well plates were covered with a breathable rayon film (catalog number 60941-086, VWR Chemicals, Radnor, Pennsylvania, USA).

For bioleaching, 150 µL of biolixiviant was transferred from each well into plates containing synthetic monazite powder covered with an adhesive polypropylene sealing Film (PCR-TS, Axygen Scientific, Union City, CA) and were incubated for 24 hours at 30 °C and 800 rpm. Final plates containing the leachate were centrifuged at 3,214 × g for 12 minutes to obtain the supernatant to be used for analysis.

For the preliminary survey, bioleaching was performed in presence of synthetic monazite mineral Mon1 (**Supplementary Data S2**) at 0.33% pulp density to facilitate an initial identification of disruption strains with differential bioleaching capabilities. For the screen with Mon2 (**Supplementary Data S2**), the pulp density was 1%. After centrifugation, a 1:10 dilution of the supernatant from each well was transferred to polystyrene flat-bottom 96-well plates filled with 100 µL of 60 µM AS-III Solution. For validation with Mon2, 100 µL of the supernatant was used instead. Analysis plates were shaken for 1 minute, then the absorbance at 650 nm was measured with the plate reader.

### REE-Bioleaching Screen Analysis

Several factors were considered to confirm the quality of the data collected, namely cross contamination of wells, systematic differences among wells caused by their position in the plate, and additional errors generated during manipulation.

To address these challenges, the raw data collected through our preliminary high-throughput survey with Mon1 was analyzed with a correcting software^75^ to detect and remove systematic errors. This software allowed us to normalize our data based on *Z*-score transformation within plates^76^. Wells with growth OD600 < 0.2 were not considered to avoid the potential offsets during the normalization process.

An initial hit identification was carried out by comparing the normalized absorbance values of the disruption mutants to the mean of the plate they were analyzed within. In a conservative manner, disruption mutants presenting changes of 1.5 standard deviation away from the plate mean were selected for further screening.

For our screen with Mon2, selected disruption mutants’ wells were picked to inoculate new 96-well plates filled with media and supplemented with antibiotic. To guarantee reliable data collection we again arrayed the mutants into a checkerboard pattern; included each mutant in triplicate; avoided perimeter wells to reduced media evaporation; and included a proxy wild-type strain within each plate.

Raw absorbance data was used for statistical analysis, which included pairwise comparisons conducted utilizing the emmeans package in R, while applying a false discovery rate (FDR) correction for *p*-value adjustment for multiple comparisons^77^. Disruption mutants that demonstrated significant differences in REE extraction were considered promising.

### Gene Ontology Enrichment Analysis

A gene ontology (GO) enrichment analysis was carried out as described by Schmitz *et al*.^16^. Enrichment was performed with the BioConductor topGO package^45,78^, employing the TopGO Fisher test with the default weight algorithm, applying a significance threshold of 0.05.

### Direct REE-Bioleaching Measurements

Selected disruption mutants that included five significant hits identified through the REE-bioleaching screen and other seven additional disruption mutants were used for further validation through bench-scale bioleaching experiments^79^ with Mon2. Additional disruption mutants were picked based on the features disrupted that represented a trait of interest, these included specific enzymes or disruptions with interesting behavior throughout the screen.

A total of 12 disruption mutant strains were plated on YPM and kanamycin agar (05039, VWR Chemicals, Radnor, PA) petri dishes for colony isolation. Three biological replicates for each disruption mutant were used to inoculate 3 mL cultures, as well as for wild-type controls in addition to no-bacteria control. After incubation for two nights at 30 °C and 200 rpm in a shaking incubator (29314, Infors Inc., Laurel, MD), cultures were normalized to OD590 0.01 in 50 mL flasks with a final volume of 5 mL and grown for another two nights. Following incubation, the cultures were diluted in half glucose 40% (w/v) to reach a final concentration in solution of 20%. Flasks were incubated at 30 °C and 200 rpm for 48 hours to allow biolixiviant production.

Bioleaching experiments were set up to digest Mon2 at 1% pulp density, for 24 hours at 30 °C and 200 rpm. Once obtained, the leachate was filtered through a 0.45 µm AcroPrep Advance 96-well filter plate (8029, Pall Corporation, Show Low, AZ, USA) by centrifugation at 1,500 × g for 5 minutes.

Each sample underwent a 20-fold dilution using 2% trace metal grade nitric acid (JT9368, J.T. Baker, Radnor, Pennsylvania, USA). Neodymium concentration in each sample was measured using an Agilent 7800 ICP-MS. Calibration and verification utilized a REE mix standard (67349, Sigma-Aldrich, St. Louis, Missouri, USA) and a rhodium internal standard (catalog number 04736, Sigma-Aldrich, *m*/*Z* = 103). Rigorous quality checks included intermittent assessment of standards, blanks, and replicate samples^80^. Data interpretation was carried out using MassHunter software, version 4.5.

A proxy wild-type strain (pWT) was selected for comparison to account for changes in the wild-type background introduced by the presence of the antibiotic cassette in the genome. The selected pWT contains a transposon insertion in an intergenic region (at base pair 1,262,471; between genes *GO_2147* and *GO_2148*) and does not demonstrate any notable phenotypic changes. This strain can be located in plate 130, well A2 in the progenitor collection.

For each disruption mutant, bioleaching measurements (*n* = 3) were evaluated against those of the wild-type and proxy wild-type strains (*n* = 3). This was done performing pairwise comparisons with Bonferroni corrected *t*-tests using the emmeans package in R. The total REE extracted was deemed significantly different if *p* < 0.05/*N* (*N* = 12).

### Limitations of the Study

During high-throughput screening, growth hindrance due to strain disruption poses considerable challenges. For example, despite the *pstS* gene disruption being flagged as a key player in REE-bioleaching during preliminary testing, its pronounced effect on microbial growth required its exclusion from final analyses because it did not meet the established growth threshold for inclusion.

Future studies could address these growth-related issues by incorporating an up-regulation collection into the screening process. This approach would not only enhance the insights gained from knockout collections but also allow for the potential identification of essential genes as significant contributors to bioleaching. Furthermore, confirmation of gene involvement of hits identified in this study will require further experiments, including gene deletion and complementation, to fully evaluate gene function and interaction.

While our study adds to the growing body of work on microbial metal recovery, the application of engineered microbes in bioleaching is still emerging. Successes have been noted with engineered strains in controlled settings such as stirred-tank reactors. Nonetheless, advancing this technology will require careful consideration of containment strategies and regulatory adjustments to appropriately mitigate potential risks.

### Statistics and Reproducibility

In our genome-scale screen, statistical robustness was ensured through multiple replicates and a rigorous validation approach. Our screen included a preliminary survey carried out in duplicate, followed by a validation screen performed in triplicate. This multistep approach allowed for the establishment of robustness in our findings. To control for potential biases, we considered factors such as cross-contamination of wells, position-related systematic noise, and errors during manipulation.

The data from our preliminary survey with Mon1 was corrected using specialized software to remove systematic errors^75^, and *Z*-score transformation was applied within plates for normalization. Wells that showed growth OD600 less than 0.2 were excluded to avoid potential offsets during this normalization process. Disruption mutants exhibiting changes of 1.5 standard deviations from the plate mean were selected for further screening.

In the subsequent validation screen with Mon2, disruption mutants were subjected to a more stringent validation process. This included a balanced distribution of mutants across the plates, triplicate data collection, and within-plate comparison strains. Statistical significance for the REE extraction capabilities of each mutant was determined using pairwise comparisons with the emmeans package in R and applying a false discovery rate (FDR) correction for *p*-value adjustment in multiple comparisons.

Bench-scale bioleaching experiments further confirmed the reproducibility of our findings, with each mutant strain’s performance benchmarked against that of the wild-type and the proxy wild-type strains. Neodymium extraction levels measured by ICP-MS were compared pairwise, with significance established at *p* < 0.05 after Bonferroni correction (*N* = 12). These stringent statistical analyses ensured that the results presented are both robust and reproducible, providing a reliable foundation for future explorations into the genetic determinants of REE-bioleaching.

