## Supplementary material for "Direct Genome-Scale Screening of *Gluconobacter oxydans* B58 for Rare Earth Element Bioleaching": Supplementary Information.pdf

### Supplementary Information Figures

**Figure S1.** Comparison of bioleaching by identified gene disruption mutants with *G. oxydans* wild-type.

### Supplementary Information Datasets

**Supplementary Data S1.** Catalog of *G. oxydans* B58 Quality-Controlled whole genome knockout collection.

**Supplementary Data S2.** Characterization of synthetic monazite powders.

**Supplementary Data S3.** Absorbance Data collected through preliminary survey of *G. oxydans* B58 Quality-Controlled Whole Genome Knockout Collection with Mon1.

**Supplementary Data S4.** *Z*-scores of gene disruptions identified through the genome-scale screen with Arsenazo-III REE-chelating dye not previously implicated in bioleaching.

**Supplementary Data S5.** Raw absorbance data for hit validation screen of REE-bioleaching from Mon2.

**Supplementary Data S6.** Gene ontology analysis.

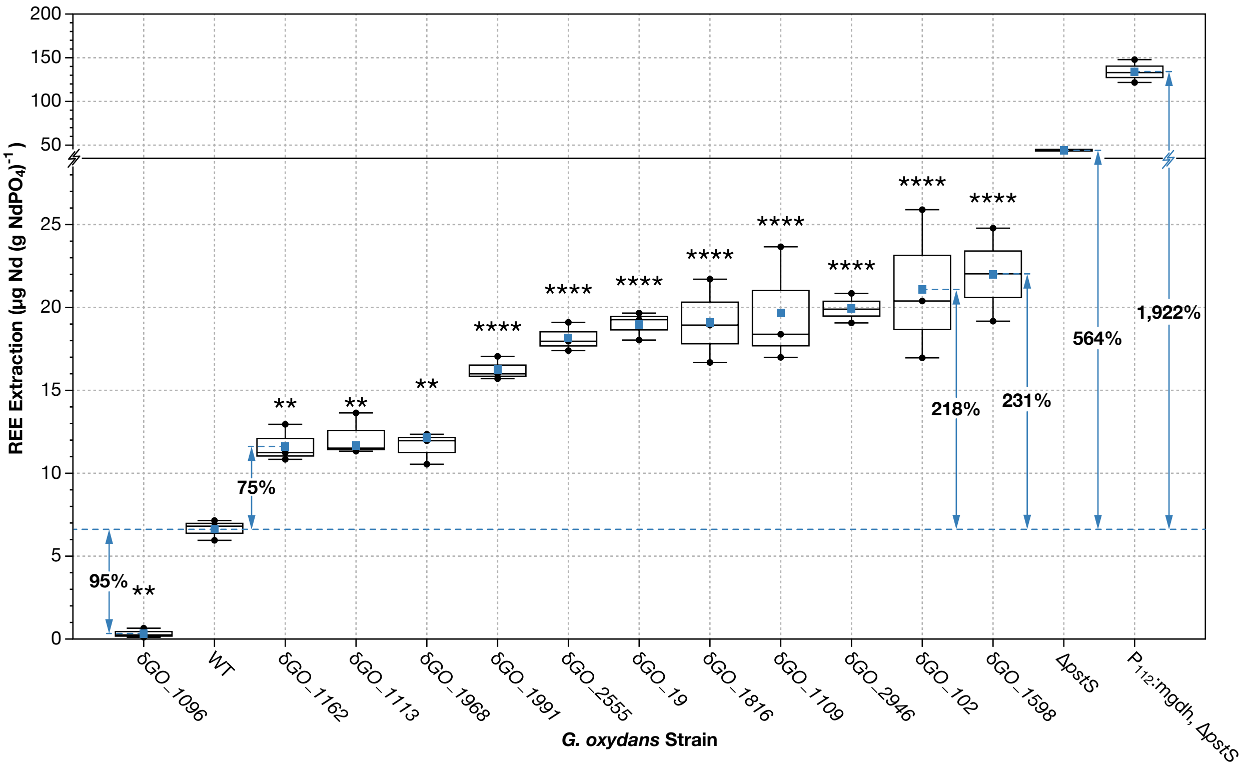


**Figure S1. Comparison of bioleaching by identified gene disruption mutants with *G. oxydans* wild-type.** Direct measurement of bioleaching extraction was performed on disruption mutants of interest after bioleaching experiments with synthetic monazite using ICP-MS analysis. REE extractions were analyzed with pairwise comparisons among disruption mutants ($n$ = 3) and the wild-type strain ($n$ = 3). Extraction levels significantly different from *G. oxydans* wild-type are labeled with asterisks (* $p < 0.05$; ** $p < 0.01$; *** $p < 0.001$; **** $p < 0.0001$) and represent statistical significance after Bonferroni correction ($N = 12$). Blue squares indicate the mean extraction for each mutant, the center line denotes the median, boxes show the upper and lower quartiles, and whiskers extend to the range of data points within 1.5 times the interquartile range. All strains analyzed were statistically significant. Disruption mutant δ*GO_1096* demonstrated a 95% reduction in extraction, while all other disruption mutants showed increased extraction.
